# Hitching a ride in the phyllosphere: Surfactant production of *Pseudomonas* spp. causes co-swarming of *Pantoea eucalypti* 299R

**DOI:** 10.1101/2023.11.07.566084

**Authors:** Michael Kunzler, Rudolf O. Schlechter, Lukas Schreiber, Mitja N.P. Remus-Emsermann

## Abstract

Here we demonstrate the beneficial effect of surfactant-producing Pseudomonads on the phyllosphere model strain *Pantoea eucalypti* 299R. To do so, we conducted a series of experiments in environments of increasing complexity. *P. eucalypti* 299R and *Pseudomonas* sp. FF1 (Pff1) or *P. eucalypti* 299R and the surfactant-deficient mutant *P*. sp. FF1::ΔviscB (Pff1ΔviscB) were co-inoculated in broth, on swarming agar plates, and on plants. In broth, there were no differences in the growth dynamics of *P. eucalypti* 299R when growing in the presence of Pff1 or Pff1ΔviscB. By contrast, on swarming agar plates, *P. eucalypti* was able to co-swarm with Pff1. Co-swarming led to a significant increase in the area colonized and, consequently, a boost in total biomass when compared to *P. eucalypti* growing with Pff1ΔviscB or in monoculture. Finally *in planta*, there were no significant effects on the population density of *P. eucalypti* 299R during co-colonization of *Arabidopsis thaliana*. However, utilizing the single cell bioreporter for reproductive success (CUSPER), we found a temporally distinct beneficial effect of Pff1 on co-inoculated *P. eucalypti* 299R subpopulations that did not occur in presence of Pff1ΔviscB. This led us to formulate a model for the positive effect of surfactant production during leaf colonization. To generalize our results, we tested the effect of three additional surfactant-producing Pseudomonads and their respective surfactant knockout mutants on *P. eucalypti* 299R on swarming agar. Resulting in similar co-swarming patterns in *P. eucalypti* 299R and showing that this organism is able to take advantage of competitors during surface colonization. Our results indicate that surfactant-dependent co-motility might be common during leaf colonization and add yet another facet to the already manyfold roles of surfactants.

## Introduction

The microbial habitat that is presented by leaf surfaces, the so-called phyllosphere, is densely colonized by microorganisms. Among these microorganisms, bacteria are the most abundant and prevalent group, establishing non-random patterns of colonization on leaves. Bacteria have a tendency to aggregate with one another in close proximity, rather than being distributed homogeneously along the leaf surface, suggesting the role of deterministic processes on leaf colonization (R. O. Schlechter, Miebach, and Remus-Emsermann 2019). In recent years, much attention has been paid to leaf colonizers and the factors that drive leaf colonization at the population and the community level. For example, the host plant species influences leaf colonization and bacterial community composition (Mercier and Lindow 2000; Bodenhausen et al. 2014; Lajoie and Kembel 2021), leading to recurring patterns of bacterial communities on leaves, at least at low phylogenetic resolution, from year to year (Vorholt 2012; Howe et al. 2023).

At the same time, microbe-microbe interactions affect community assemblages on leaves (Carlström et al. 2019; Martin Schäfer et al. 2023; M. Schäfer, Vogel, and Bortfeld-Miller 2022). However, many of the mechanisms driving these interactions remain unclear. In studies where bacterial communities are relatively simple, it appears that nutrient overlap has a limited impact on leaves (R. O. Schlechter et al. 2023; Remus-Emsermann and Schlechter 2018; Martin Schäfer et al. 2023). This suggests that the highly segregated leaf environment constrains interspecies bacterial interactions, despite their tendency to co-aggregate (Monier and Lindow 2005; Remus-Emsermann et al. 2014). As proximity is a key factor for interaction, mobility might emerge as an important factor that is shaping interactions and, ultimately, bacterial communities. In general, mobility on leaves has received limited attention, with most studies being related to the presence of flagellar genes in *Pseudomonas* spp. (van der Wal et al. 2013; Haefele and Lindow 1987; Delmotte et al. 2009). Since roughly 5% of the leaf surface is colonized by bacteria (Remus-Emsermann et al. 2014) and not all leaf-colonizing bacteria are flagellated (Bai et al. 2015), other means of mobility must be present on leaves. This may include movement mediated by colony expansion or gliding motility (Su et al. 2012; Martínez, Torello, and Kolter 1999), or spread by dew (Van Stan et al. 2020; Beattie 2011). The genera *Pantoea* and *Pseudomonas* are known to be a very common leaf colonizer (Rastogi et al. 2012; Vorholt 2012). Both taxa interact with their plant host in various ways, ranging from pathogenicity to plant protection (Xin, Kvitko, and He 2018; Zengerer et al. 2018; Ramette et al. 2011; Vrancken et al. 2013).

However, their interaction within communities has not received much attention, despite some initial efforts (Monier and Lindow 2005).

Phyllosphere-colonizing Pseudomonads have previously been studied with regard to their ability to produce surfactants (S. Oso et al. 2021; Burch et al. 2014; Hernandez and Lindow 2019; Simisola Oso et al. 2019; Schreiber et al. 2005). Interestingly, a fitness advantage of surfactant-producing Pseudomonads over their surfactant-deficient counterparts was evident only under specific conditions such as fluctuating humidity (Burch et al. 2014). Otherwise, surfactant production did not seem to affect the ability of Pseudomonads to colonize leaves (S. Oso et al. 2021).

Surfactants have been suggested to have multiple roles in the phyllosphere, including increased spread of water by reducing surface tension, improved nutrient diffusion from the apoplast to the phyllosphere by increasing cuticle permeability (Schreiber et al. 2005), and increased drought resistance due to their hygroscopic nature (Burch et al. 2014; Hernandez and Lindow 2019). On semi-solid media, surfactants are necessary for swarming, and it has been shown that it can facilitate the mobilization of some bacteria in a process called co-swarming (Morin et al. 2022). However, the effect of surfactant-producing bacteria on a second colonizer in the phyllosphere has, to our knowledge, not been studied.

Bacterial mobility relies on various mechanisms. Swimming motility is an active form of movement, powered by rotating flagella, occurring in liquid or low-viscosity conditions (Ha, Kuchma, and O’Toole 2014). Gliding motility is movement on surfaces, independent of flagella or pili (McBride 2001). Another surface-associated motility mode is sliding motility, driven by excreted biosurfactants, preventing cells from forming thick biofilm layers and aiding dispersal on the surface they inhabit (Wadhwa and Berg 2022). Twitching motility is mediated by type IV pili, which propels bacteria by retraction, while swarming motility is the coordinated movement of multiple bacteria across solid or semisolid surfaces, requiring flagella, a functional quorum sensing system, and surfactant biosynthesis (Merz, So, and Sheetz 2000). With this mechanism, bacteria co-migrate in side-by-side groups called rafts, instead of individually as in swimming motility (Kearns 2010).

Previously, we developed a bioreporter, CUSPER, to measure the number of divisions of individual cells of the bacterium *Pantoea eucalypti* 299R (syn. *Erwinia herbicola* 299R, syn. *Pantoea agglomerans* 299R) after they arrive in new environments. This bioreporter for reproductive success is based on the dilution of green fluorescent protein (GFP) during cell division. Thereby, the GFP intensity becomes a direct proxy for the number of experienced divisions (Remus-Emsermann and Leveau 2010; Remus-Emsermann et al. 2012). In a previous study, we observed that *Pseudomonas* spp. increased the single-cell reproductive success of *P. eucalypti* 299R (Pe299R) *in planta* despite being strong competitors *in vitro* (R. O. Schlechter et al. 2023). Given the common feature of surfactant production in Pseudomonads (Raaijmakers et al. 2010; Simisola Oso et al. 2019), we hypothesized that surfactant production increases the reproductive success of a second colonizer in the phyllosphere. To test this, we investigated the interactions between Pe299R, surfactant-producing *Pseudomonas* isolates, and their respective knockout mutants in various conditions, ranging from liquid medium, agar surfaces, and leaves. We observed contrasting results in liquid cultures compared to agar surfaces and leaves.

## Material and methods

### Bacteria, strain construction and growth conditions

All bacteria used in this study are listed in Table 1. Bacteria were routinely grown on lysogeny broth agar (LB-Agar, HiMedia). *Pantoea eucalypti* 299R was equipped with a constitutively expressed red fluorescent mScarlet-I protein gene and the plasmid pCUSPER (R. O. Schlechter et al. 2023). The resulting strain will be referred to as Pe299R_CUSPER_ from here onwards. To maintain the plasmid carrying the reproductive success construct in Pe299R_CUSPER_, the agar was supplemented with 50 mg L^-1^ kanamycin (Roth). Constitutively cyan-fluorescent derivatives of strains *Pseudomonas* sp. Pff1 (Pff1_cyan_) and *Pseudomonas* sp. Pff1::ezTn5-viscB (Pff1ΔviscB_cyan_) used in this study were generated using plasmid pMRE-Tn7-141 (R. O. Schlechter et al. 2018) explained elsewhere using conjugation and the auxotrophic donor strain *E. coli* ST18 (R. Schlechter and Remus-Emsermann 2019; Thoma and Schobert 2009). Strains *Pseudomonas* sp. Pff2, *Pseudomonas* sp. Pff2::ezTn5-viscB, *Pseudomonas* sp. Pff3, *Pseudomonas* sp. Pff3::ezTn5-massB, *Pseudomonas* sp. Pff4, and *Pseudomonas* sp. Pff4::ezTn5-massB were characterized elsewhere and carry Tn*5-*knockout insertions in the viscosin B (*viscB*) or the massetolide B (*massB*) genes, respectively (S. Oso et al. 2021). Those genes were previously shown to be responsible for surfactant production.

**Table 1.** Bacterial strains used in this study.

| Name | Relevant genotype/<br>properties | Antibiotic<br>resistance | Abbreviation | Origin |
| --- | --- | --- | --- | --- |
| *Escherichia coli* ST18 | pMRE-Tn7-145;<br>red fluorescent | Cm, Gent,<br>Amp |  | (R. O. Schlechter et al. 2018) |
| *Escherichia coli* ST18 | pMRE-Tn7-141;<br>cyan fluorescent | Cm, Gent,<br>Amp |  | (R. O. Schlechter et al. 2018) |
| *Pantoea eucalypti* 299R |  | Rif |  | (Remus-Emse rmann et al. 2013) |
| *Pantoea eucalypti* 299R | ::Tn7-145; pCUSPER;<br>red fluorescent, growth<br>bioreporter | Rif, Gent,<br>Kan | Pe299R<sub>CUSPER</sub> | (R. O. Schlechter et al. 2023) |
| *Pseudomonas* sp. FF1 |  |  | Pff1 | (S. Oso et al. 2021) |
| *Pseudomonas* sp. FF1 | ::Tn7-141;<br>cyan fluorescent | Gent | Pff1<sub>cyan</sub> | This study |
| *Pseudomonas* sp. FF1 | ::ezTn5-viscB | Kan | Pff1ΔviscB | (S. Oso et al. 2021) |
| *Pseudomonas* sp. FF1 | ::ezTn5-viscB::Tn7-141;<br>cyan fluorescent | Gent, Kan | Pff1ΔviscB<sub>cyan</sub> | This study |
| *Pseudomonas* sp. FF2 |  |  | Pff2 | (S. Oso et al. 2021) |
| *Pseudomonas* sp. FF2 | ::ezTn5-viscB | Kan | Pff2ΔviscB | (S. Oso et al. 2021) |
| *Pseudomonas* sp. FF3 |  |  | Pff3 | (S. Oso et al. 2021) |
| *Pseudomonas* sp. FF3 | ::ezTn5-viscB | Kan | Pff3ΔmassB | (S. Oso et al. 2021) |
| *Pseudomonas* sp. FF4 |  |  | Pff4 | (S. Oso et al. 2021) |
| *Pseudomonas* sp. FF4 | ::ezTn5-viscB | Kan | Pff4ΔmassB | (S. Oso et al. 2021) |

#### *In vitro* co-inoculation assay

To investigate the co-colonization of Pe299R_CUSPER_, Pff1_cyan_ or Pff1ΔviscB_cyan_ *in vitro*, all strains were grown in 3 mL M9 minimal media (Na_2_HPO_4_.7H_2_O 64 g L^-1^, KH_2_PO_4_ 15 g L^-1^, NaCl 2.5 g L^-1^, NH_4_Cl 5.0 g L^-1^) supplemented with 200 μM FeCl_3_ (to avoid production of autofluorescent pyoverdines by Pseudomonads) and 0.13% w/v glucose, fructose and sorbitol each (M9 3C) overnight at 30°C and 200 rpm. Five hundred μL of each overnight culture were then used to inoculate 50 mL M9 3C, respectively. After six hours, cultures were washed twice by centrifugation at 2,000 *g* for 5 minutes and resuspension in PBS. Suspensions were diluted to an optical density (OD600nm) of 1 and the following treatments were prepared: Pe299R_CUSPER_, Pe299R_CUSPER_ vs. Pff1_cyan_, Pe299R_CUSPER_ vs. Pff1ΔviscB_cyan_, Pff1_cyan_, and Pff1ΔviscB_cyan_. These suspensions were diluted into either M9 supplemented with glucose 0.4% w/v (M9gluc), M9 3C, or LB to a final OD600nm of 0.04 for each strain. Triplicate treatments were pipetted into a 96-well plate and placed into a CLARIOstar Plus microplate reader (BMG Labtech). The fluorescence of the mScarlet-I expressing Pe299R_CUSPER_ was determined by excitation at 530-570 nm and measuring emission at 580-620 nm while *Pseudomonas* produced mTurquoise2 was excited at 400-440 nm and its emission was measured at 450-490 nm using the CLARIOstar software (version 5.70, BMG Labtech). Measurements were performed in 20-min intervals for 42 hours. The plate was incubated at 30°C and the plate was shaken in a double orbital at 200 rpm between measurements.

The area under the curve of the red and cyan fluorescence kinetic of each sample was determined in Prism 10.0.1 (Graphpad). As cyan fluorescence kinetics exhibited a negative change in fluorescence that was media dependent, the data was corrected by a constant value as to move every datapoint above the baseline before the absolute area under the curve was determined.

### Swarming assays

To determine the swarming ability of Pe299R_CUSPER_, Pff1_cyan_, and Pff1ΔviscB_cyan_, each strain was inoculated onto soft agar. Soft agar plates were prepared with 0.83% w/v lysogeny broth agar, (LB-Agar, HiMedia) with a final agar concentration of 0.5% w/v. The center of the plate was inoculate with 10 μL of bacterial suspension (OD600nm = 1, prepared as above) and pictures were taken after 24 h of incubation at 30°C using a darfield illuminator (Parkinson 2007) in a dark chamber (MultiImage Light Cabinet) with attached Axiocam 105 (Zeiss) and Zen Core (version 3.2, Zeiss).

LB soft agar was prepared by diluting standard LB agar with ddH_2_O in a 1:2 ratio, supplemented with 200 μM FeCl_3_. Two mL soft agar was then distributed into a 6-well microtiter plate (Greiner). After the agar cured, a 1.5 μL drop of washed bacterial suspensions (adjusted OD600nm = 0.5) was placed in the middle of each well. The following monocultures and mixtures were prepared: (i) Pe299R_CUSPER_, (ii) Pe299R_CUSPER_ vs. Pff1_cyan_, (iii) Pe299R_CUSPER_ vs. Pff1ΔviscB_cyan_, and (iv) Pff1_cyan_, and Pff1ΔviscB_cyan_. The plates were then incubated at 30°C for 16 hours. Afterwards, the plates were placed into a CLARIOstar Plus Microplate reader (BMG Labtech). The fluorescence of the mScarlet-I expressing Pe299R_CUSPER_ was determined by exciting the sample at 530-570 nm and measuring emission at 580-620 nm while *Pseudomonas* produced mTurquoise2 was excited at 400-440 nm and its emission was measured at 450-490 nm. The plates were scanned using the 30 x 30 Matrix scan mode of the CLARIOstar software using bottom optics to obtain the spatial information of each strain. This resulted in a two-dimensional distribution of red and cyan fluorescence data. Background subtraction was performed individually for each datapoint. Data was visualized using the heatmap function of Prism. Total fluorescence of the red fluorescence channel was used as a proxy to determine the total biomass of Pe299R_CUSPER_. The experiment was performed four times on LB soft agar.

Additionally, to test if co-swarming also occurs when Pe299R_CUSPER_ co-colonises agar surfaces with other swarming pseudomonads, Pe299R_CUSPER_ was co-inoculated with Pff2, Pff3, and Pff4, as well as their respective surfactant deficient mutants on soft LB agar.

### Plant growth

Plants were prepared as explained in Miebach et al. (Miebach et al. 2020). Briefly, *Arabidopsis thaliana* Col-0 seeds were sterilized by vortexing in 70% v/v ethanol for two minutes. The ethanol was then removed by pipetting, and the seeds were treated with 50% v/v household bleach (NaOCl, 2.47%) and 0.02% v/v Tween-20 for 7 minutes. Afterwards, seeds were washed three times with sterile water. Sterilized seeds were kept in water and stratified in dark at 4°C for three days. After stratification, seeds were sown on cut pipette tips (∼5 mm in length) pre-filled with ½ strength Murashige-Skoog (½ MS) agar medium, placed in a petri dish with ½ MS agar. Petri dishes were closed with parafilm (Bemis) and were placed in a M-5-Z growth cabinet (PolyKlima) with a 11/13 day/night interval (including 30 minutes dusk and 30 minutes dawn during which the lights slowly increase in intensity), 80% relative humidity, and 210 μmol s^-1^m^-2^ light intensity.

Gnotobiotic culture boxes were prepared following the methods described by (Miebach et al. 2020). In brief, Magenta Culture Boxes GA-7 were filled with 90 g finely zeolite clay granulate (Klinoptilolith, 0.2-0.5 mm, Labradorit.de) and autoclaved with lids closed. Lids were previously perforated and holes were covered with a double layer of gas permeable tape (Micropore, 3M). After autoclaving and cooling down, 45 ml of sterile ¾ MS Medium was added per box under aseptic conditions. One week after sowing, seedlings were transferred, including the tips they were germinated on, into prepared Magenta boxes. Four seedlings were placed per box and grown under the same conditions described above.

### Plant inoculation

To prepare the inoculum of the different strains, a single colony was selected and used to prepare overnight cultures in LB medium (HiMedia, *Pe*299RCUSPER with 50 mg L^-1^ kanamycin respectively). The overnight cultures were then used to inoculate fresh liquid cultures by adding 500 μl of cultures to 50 ml fresh medium. Those cultures were grown to mid exponential phase while shaking at 30°C and 200 rpm. *Pe*299R_CUSPER_ cultures were supplemented with kanamycin and 1 mM isopropylthio-β-galactoside (IPTG) to induce GFP-expression (Remus-Emsermann and Leveau 2010). When reaching log-phase after approximately 6 hours of growth, cells were pelleted by 5 minutes of centrifugation at 4000 × *g*, washed three times and adjusted to an OD600nm of 0.1 with sterile phosphate buffered saline (PBS, 1.37 M NaCl, 27 mM KCl, 100 mM Na_2_HPO_4_, 18 mM KH_2_PO_4_, pH 7). Where appropriate, strains were mixed prior inoculation, resulting in a relative OD600nm = 0.05 for each strain.

To inoculate *A. thaliana* plants, Magenta® box lids were temporarily replaced with a lid with only one central hole. Each box was then sprayed with 200 μl of inoculum with an airbrush paint gun (Ultra Spray gun, Harder & Steenbeck, Norderstedt, Germany). Then, the lids were replaced and plants were placed back into the growth chamber.

### Plant sampling and bacterial cell recovery

Three-week-old plants were sampled after 0 h, 12 h, 18 h and 24 h post inoculation. Plants were sampled by harvesting the total aboveground material using sterile forceps and scissors. Plant material was transferred into a 15-ml centrifuge tube, and fresh weight was determined before 1 ml PBS was added to recover leaf surface-attached bacteria. The samples were then vortexed for 15 seconds and sonicated for 5 minutes at 75% intensity in a sonication bath (Emmi 12 HC, EMAG). Leaf washes were recovered from the tubes and a 100 μl aliquot was used to determine colony forming units (CFU) by serial dilution on LB agar supplemented with rifampicin (50 mg/L) and kanamycin (50 mg/L) to select for Pe299R_CUSPER_. The remaining leaf wash was centrifuged for 10 minutes at 15,000 x *g* at 4°C, the supernatant was discarded, and the resulting bacterial pellet resuspended was fixed overnight in 4% w/v paraformaldehyde in PBS at 4°C. After fixation, the PFA was removed by three washing steps with sterile PBS. After the last washing step, the pellets were resuspended in 50 μl PBS mixed with 50 μl 96% v/v ethanol and stored at -20°C until they were analyzed. All samples were analyzed within two weeks.

### Microscopical analysis of CUSPER signals and image cytometry

Recovered bacterial cells were analyzed using widefield fluorescence microscopy as described elsewhere (R. O. Schlechter et al. 2023). Briefly, cells were drop spotted onto 1% w/v agarose slabs on a microscopy slide and covered with a cover slip. An AxioImager Z2 microscope (Zeiss) with a Axiocam 712 mono camera (Zeiss) and X-cite Xylis broad spectrum LED light source (Excelitas) were used. Images were acquired at 1000× magnification (EC Plan-Neofluar 100×/1.30 Ph3 Oil M27 objective) in phase contrast, green fluorescence and red fluorescence (Zeiss filter sets 38HE (BP 470/40-FT 495-BP 525/50) and 43HE (BP 550/25-FT 570-BP 605/70), respectively) using the software Zen 3.3 (Zeiss). At least 120 cells were acquired per biological replicate which consisted of bacteria pooled from four different plants. Images were analyzed using FIJI and as described previously (Schindelin et al. 2012; R. O. Schlechter et al. 2023). Briefly, Pe299_CUSPER_ cells were identified using their constitutive red fluorescence and the thresholding method “intermodes”. The resulting regions of interest were converted into a binary mask. Artifacts were excluded by analyzing only particles sizes from 0.5–2.5 μm and excluding cells touching the image edges. All images were manually curated using the phase contrast images to exclude false positive red fluorescent particles. The mask was then used to determine green fluorescence intensity of Pe299R_CUSPER_ cells. In addition, the average background fluorescence was measured by randomly sampling the background area of each image. Background fluorescence was subtracted from the data. As previously described, the number of experienced divisions of every cell was determined by calculating log_2_(average cell’s GFP fluorescence at t=0 divided by single cell’s fluorescence at time t) (Remus-Emsermann and Leveau 2010).

## Results

### Pseudomonads negatively affect growth of Pe299R_CUSPER_ *in vitro*

To investigate the interaction of Pe299R_CUSPER_ with Pff1_cyan_ or Pff1ΔviscB_cyan_ in homogeneous conditions, they were grown in shaken liquid cultures. Under these conditions, there was a strong decrease of Pe299R_CUSPER_ red fluorescence when it was co-inoculated with Pff1_cyan_ or Pff1ΔviscB_cyan_ (Figure 1). Different media had slightly different effects on this interaction and the decrease ranged from >90% decrease in M9gluc or M9 3C to >30% reduction in LB. This effect was not associated with the *Pseudomonas* red autofluorescence. By contrast, the effect of Pe299R_CUSPER_ on Pff1_cyan_ and Pff1ΔviscB_cyan_ was much smaller. Only in rich LB growth medium, we could detect a significant negative effect of Pe299R_CUSPER_ on the Pseudomonads (Supplemental Figure 1). In conclusion, the Pseudomonads had a strong negative effect on Pe299R_CUSPER_ *in vitro*, while Pe299R_CUSPER_ is barely affecting the growth of the Pseudomonad strains.

**Figure 1.**
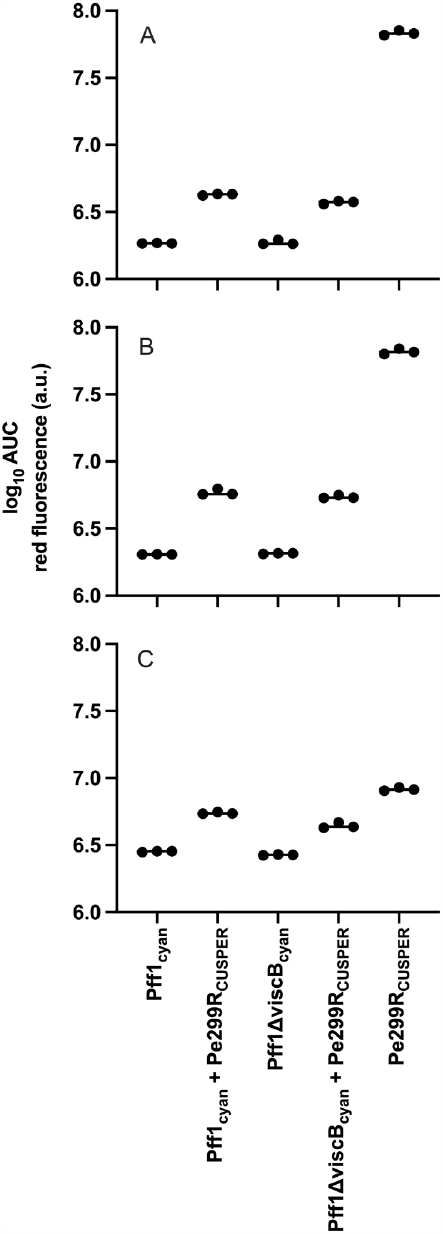
Impact of Pff1_cyan_ and Pff1ΔviscB_cyan_ on the growth of Pe299R_CUSPER_ in different media. The effect on Pe299R_CUSPER_ growth was measured by determining the area under the curve of the red fluorescence of Pe299R_CUSPER_ monocultures and co-inoculations of Pe299R_CUSPER_ and the Pseudomonads. A) M9 supplemented with glucose B) M9 supplemented with glucose, fructose and sorbitol and C) LB. Note that the y-axis is on a log_10_ scale.

### Swarming on solid media

When inoculated onto soft KB agar media alone, the different strains showed various swarming patterns. Pe299R_CUSPER_ alone formed as a colony with entire edges that is slightly more slimy compared to growth on standard agar media and does not grow beyond the initial location of inoculation. Pff1_cyan_ grew in a flower-shaped colony that is indicative for swarming, far beyond the initial site of inoculation. By contrast, Pff1ΔviscB_cyan_ formed a colony similar to Pe299R_CUSPER_ and was not able to move far beyond the initial site of inoculation.

To investigate bacterial behavior after co-inoculating Pe299R_CUSPER_ with the two different Pseudomonads, we used their respective constitutively expressed fluorescent proteins to track their biomass on a swarming agar (Figure 3). While monocultures behaved as expected (Figure 3 A), we found that co-inoculated bacteria affected each other in different fashions. In co-culture with the swarming Pff1_cyan_, Pe299R_CUSPER_ had a much wider distribution on the agar surface (Figure 3 B). By contrast, in co-culture with the non-swarming Pff1ΔviscB_cyan_, Pe299R_CUSPER_ was similarly restricted in its distribution as in a monoculture. By repeating the experiment four times, we were able to determine the total change in biomass during mono and co-cultures (Figure 3 C). As a result, the Pe299R_CUSPER_ biomass, as measured by red fluorescence, was significantly higher in co-inoculations with surfactant producing Pff1_cyan_ compared to Pe299R_CUSPER_ monocultures and co-inoculations with non-surfactant producing Pff1ΔviscB_cyan_ (*p* = 0.0001 and <0.0001, respectively, one-way ANOVA Tukey’s multiple comparison test). By contrast, albeit not significant, Pe299R_CUSPER_ biomass is slightly reduced during co-culture with Pff1ΔviscB_cyan_.

### Co-inoculation *in planta* does not significantly affect Pe299R_CUSPER_ at the population scale but does affect reproductive success at the single cell resolution

To investigate the effect of a co-colonizer producing surfactant on Pe299R_CUSPER_ *in planta*, the strains were co-inoculated onto axenic *A. thaliana* plants. The two different co-colonizing Pseudomonads affected the population size of Pe299R_CUSPER_ similarly. Although we observed a trend that the populations were slightly higher after 24 hours in one of the experiments, this was not consistent between experiments (Supplemental figure 2).

**Figure 2.**
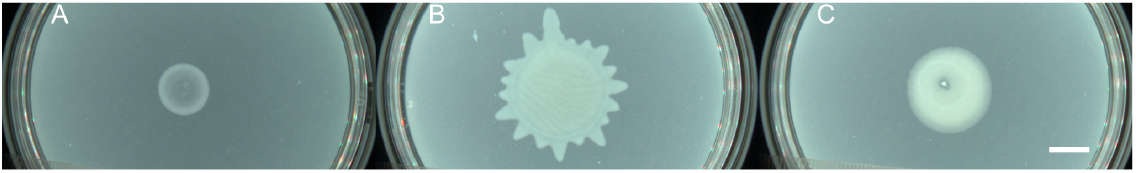
Colony morphology of A) Pe299R_CUSPER_ B) Pff1_cyan_ and C) Pff1ΔviscB_cyan_ on KB swarming agar. Pictures were taken in a darkfield illuminator after 24 hours of growth at 30 °C. Scale bar = 1 cm

**Figure 3.**
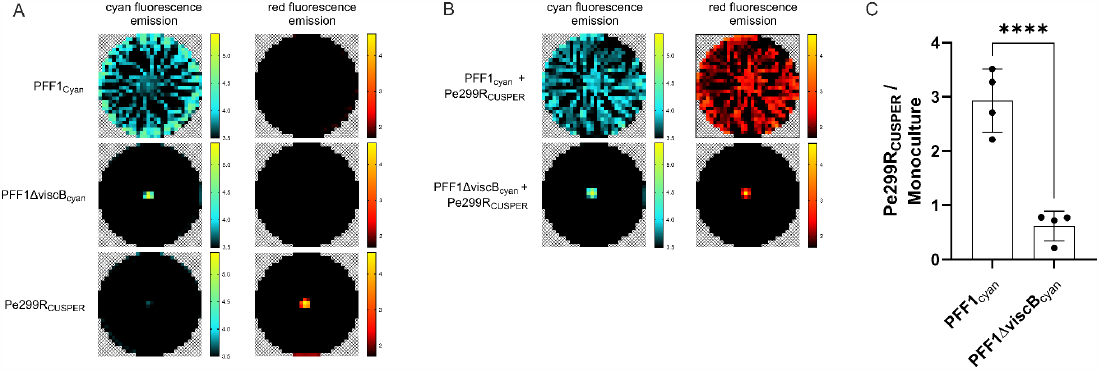
Spatially resolved analysis of bacterial growth and swarming behavior on LB soft agar plates after 16 hours. A) Monocultures of Pe299R_CUSPER_, Pff1_cyan_ and Pff1ΔviscA_cyan_ or B) mixed cultures of Pe299R::red in combination with Pff1_cyan_ or Pff1ΔviscA_cyan_ on LB soft agar plates. Bacterial growth and biomass were tracked with a fluorescent microtiter plate reader. Growth of Pe299R_CUSPER_ was determined by measuring red fluorescence emission and growth of Pseudomonads was determined by measuring cyan fluorescence emission. Note that the color scales are presented as the decadic logarithm of arbitrary fluorescence units. Furthermore, the experiment was repeated four times on LB soft agar plates. C) The fluorescence of Pe299R_CUSPER_ was determined after 16 hours and the ratio of Pe299_CUSPER_ growing in mixtures over its monoculture was determined.

By contrast, at the single cell resolution, there are noteworthy differences in the population development during the co-colonization of leaves (Figure 4). Generally, a proportionally larger subpopulation of Pe299R_CUSPER_ experienced more cell divisions in presence of the surfactant producing Pff1_cyan_ compared to the non-producing Pff1ΔviscB_cyan_. This effect is most apparent after 24 hours and was reproduced in three independent experiments (Figure 4 and Supplemental Figure 3). Furthermore, we observed that after 18h, the presence of the non-surfactant producing strain had a positive effect on Pe299R_CUSPER_ (Figure 4).

**Figure 4.**
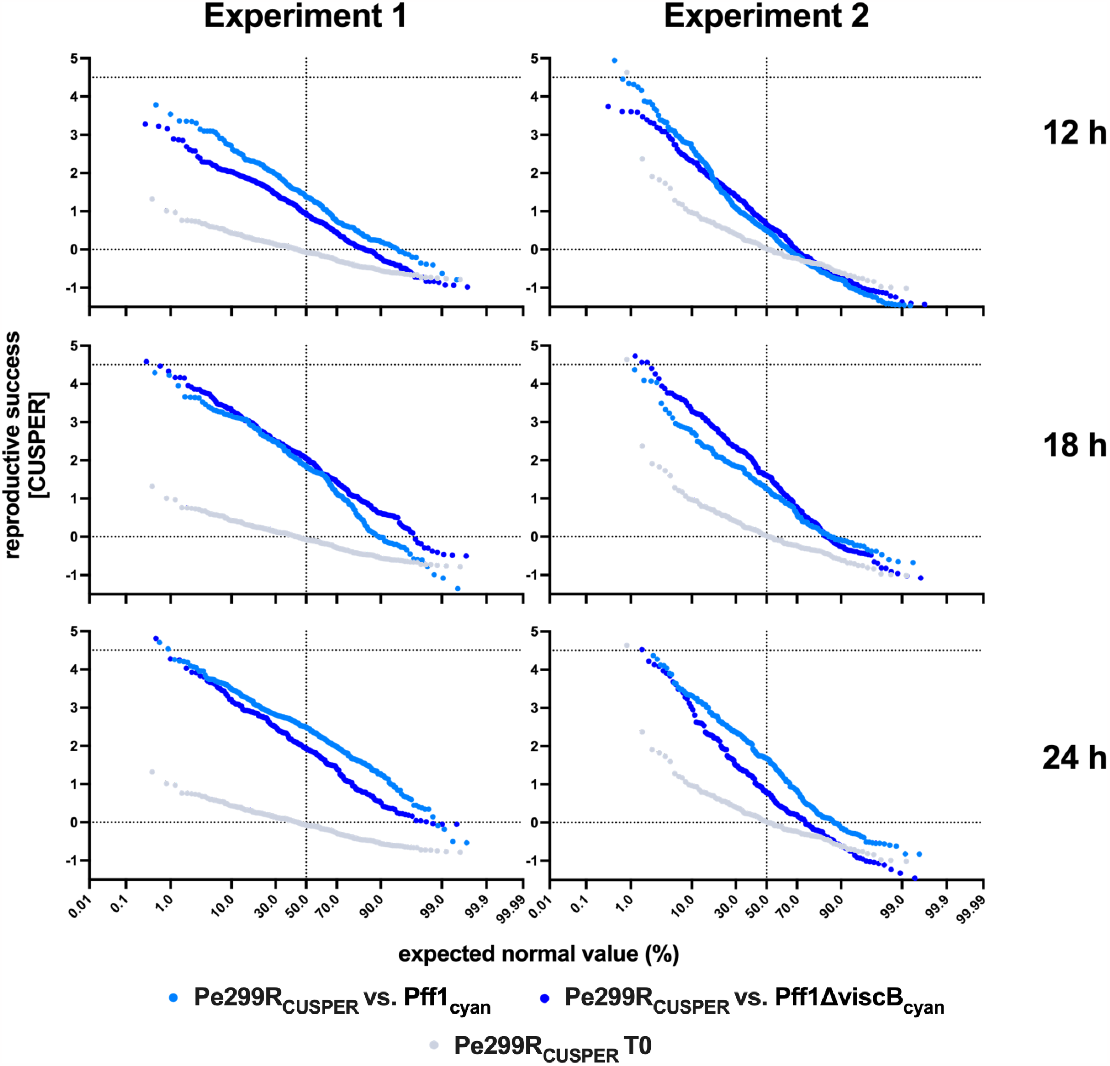
Reproductive success of individual Pe299R_CUSPER_ cells during co-colonization of leaves with Pff1_cyan_ or Pff1ΔviscB_cyan_. In gray, the respective T0 fluorescence intensity of the Pe299R_CUSPER_ population is depicted. Every increase in reproductive success depicts a cell division relative to the T0 population. Every sample is pooled from the bacteria recovered from four plants.

## Discussion

The potential roles of biosurfactants in plant-microbe interactions have been investigated from many different angles: their hygroscopic nature enhances the survival of Pseudomonads (Burch et al. 2014) and increases water availability on leaves (Hernandez and Lindow 2019), they are suspected to increase diffusion of nutrients through the hydrophobic leaf cuticle (Schreiber et al. 2005), and they have been shown to increase alkane degradation by leaf colonizing bacteria (S. Oso et al. 2021). Generally, the abundance of biosurfactant producing bacteria on leaves is proportionally high (Schreiber et al. 2005; Burch et al. 2016; Simisola Oso et al. 2019), indicating a potential ecological importance during life on leaves. Here, we show that surfactant producing bacteria may facilitate the growth of co-colonizers on leaves in a surfactant production and surface colonization dependent manner.

Being both copiotrophic generalists that exhibit a wide range of nutrient utilization, it was expected that Pe299R_CUSPER_ and both Pseudomonads affected each other strongly in liquid culture. Indeed, it was mostly Pe299R_CUSPER_ that was negatively affected in minimal media and less so in complex LB medium. In turn, the Pseudomonads were almost not affected at all by the presence of Pe299R_CUSPER_ in minimal media and to a smaller degree in LB medium. These results are in stark contrast to our observations on a spatially structured soft agar surface. Here, the non-surfactant producing Pff1ΔviscB_cyan_ negatively affected Pe299R_CUSPER_, but not Pff1_cyan_ which instead increased the amount of Pe299R_CUSPER_ fluorescence as a proxy for biomass by a factor of three and thereby increased its fitness dramatically. As this increase in correlates to a larger spread of the Pe299R_CUSPER_, it is likely that this increase can be accredited to a mobilization of Pe299R_CUSPER_ by Pff1_cyan_, similar co-swarming phenomena have previously been observed in *Paenibacillus vortex* and other *Paenibacillus* strains (Finkelshtein et al. 2015), as well as *Pseudomonas aeruginosa* and *Burkholderia cenocepacia* (Venturi et al. 2010; Morin et al. 2022). Generally, it has been shown that the swarming conferring strain gains a benefit from the co-swarmer by for instance taking advantage of antibiotic resistances provided by the immobile strain (Finkelshtein et al. 2015), or by recovering mobility in case one strain loses mobility factors (Venturi et al. 2010; Morin et al. 2022). In our study, there is no apparent benefit for the strains that mobilize Pe299R_CUSPER_. Instead, it is Pe299R_CUSPER_ that seems to be able to take advantage of the swarming strains and increases its fitness as compared to homogeneous shaken liquid cultures where swarming does not confer any fitness advantages. This observation led us to test if this effect also pertains to colonization of leaf surfaces, which is the origin of isolation of all strains used in this study.

To test this, we tracked the changes of Pe299R_CUSPER_ populations on *A. thaliana* and the ability of Pe299R_CUSPER_ individual cells to divide on leaves, exploiting the reproductive success bioreporter CUSPER. At the population scale, the effect of co-colonization was minimal, although a trend of higher Pe299R_CUSPER_ populations during co-colonization with Pff1_cyan_ as compared to co-colonization with Pff1ΔviscB_cyan_ could be observed (Supplemental Figure 2). This is in contrast to single cell observations, where we could detect an increased number of cells that experienced a notably higher number of divisions in presence of Pff1_cyan_ as compared to the presence of Pff1ΔviscB_cyan_ (Figure 4) after 24 hours. Curiously, this effect was not visible after 18 hours where Pe299R_CUSPER_ seemed to have a minimal growth advantage in presence of Pff1ΔviscB_cyan_. To explain this, we hypothesize the following model of interactions: During inoculation, it is more probable for individual strains to arrive on the leaf without a competitor in their local environment (Figure 5 A). In a scenario without swarming (Figure 5 B and C), the chance of bacterial strains interacting with each other is low as the minimal interaction distance between bacteria on leaves is about 10 μm (Esser et al. 2015; Remus-Emsermann et al. 2014). As a consequence, bacteria can initially grow rather unimpeded by competition. By contrast, in a scenario with swarming (Figure 5 D and E), the swarming strain will have higher chances to meet the non-swarming Pe299R_CUSPER_, which leads to high local competition and locally reduced populations of Pe299R_CUSPER_ (compare Figure 3, where the local biomass of Pe299R_CUSPER_ is reduced during competition). However, as co-swarming leads to mobilization of Pe299R_CUSPER_ and allows the exploration of new microenvironments, this should lead to an increased reproductive success of the strain despite the local competition (Figure 5 E). This is directly analogous to the scenario observed on the spatially structured agar surface (Figure 3).

**Figure 5.**
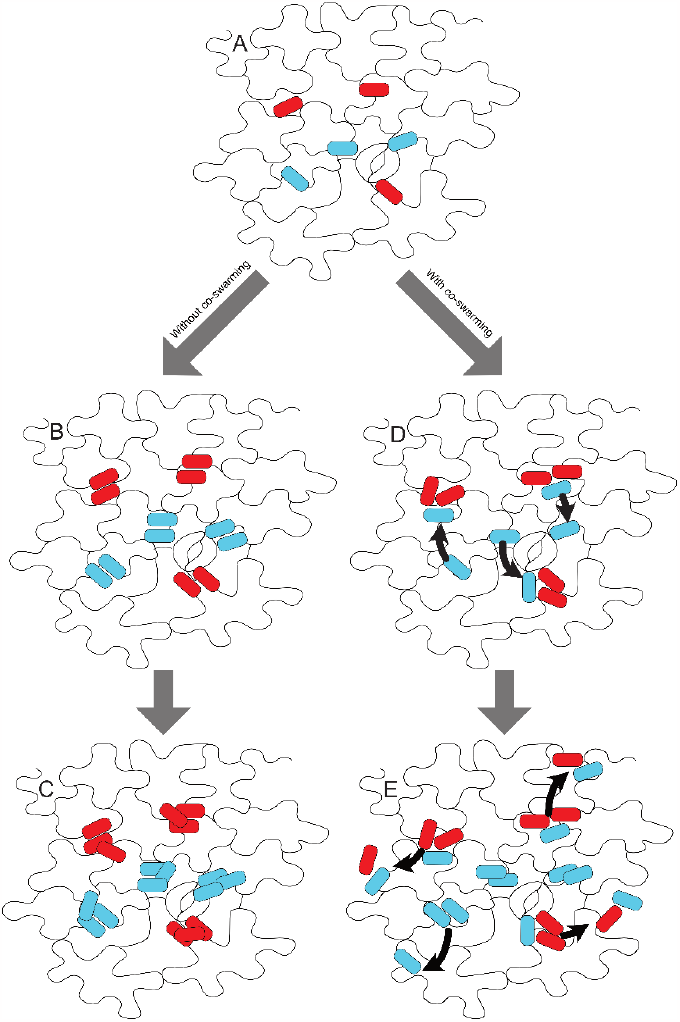
Model explaining behavior of Pe299_CUSPER_ after co-inoculation with Pff1ΔviscB_cyan_ (B, C) or Pff1_cyan_ (D, E). A) depicts the initial distribution of non-motile, red-colored Pe299_CUSPER_ cells and co-inoculated cyan-colored Pff1ΔviscB_cyan_ or Pff1_cyan_ cells. In the left scenario, cyan cells do not produce any surfactants and do not swarm. As a result red and cyan populations remain localized and rely on the locally available nutrients. In the right scenario, cyan cells produce surfactants and swarm. This leads to a higher probability of cyan cells to encounter red cells, which leads to a temporally high competition for nutrients and a reduction of red cell divisions (D). However, if red cells are mobilized by co-swarming as a results of close spatial proximity to cyan cells, this leads to the exploration of new sites and an increase of growth.

### Effect of other surfactant-producing Pseudomonads on Pe299R_CUSPER_

Interestingly, in a previous study we have observed a similar effect of increased reproductive success of Pe299R_CUSPER_ when co-colonizing plant leaves with surfactant-producing *P. koreensis* P19E3 *and P. syringae* B728a despite high overlap in resource utilization abilities (R. O. Schlechter et al. 2023). This effect can now likely be accredited to the production of surfactants and co-swarming. To further test if co-swarming of Pe299R_CUSPER_ with Pseudomonads can be generalized and accredited to surfactant production, we have tested the growth Pe299R_CUSPER_ and three additional Pseudomonads and their respective surfactant knockout mutants on soft agar. Indeed, we could observe that the Pseudomonas strains Pff2 and Pff4, but not their respective surfactant knockout mutants to co-swarming of Pe299R_CUSPER_ (Supplemental Figure 4). Intriguingly, Pff3 was not able to mobilize Pe299R_CUSPER_. Pff3 also exhibited a different style of swarming on soft agar plates which seemed to consist of a very thin layer of biomass (Supplemental Figure 5), which might explain this difference in co-swarming. Exploring the reasons for those differences are beyond the scope of this study and might be growth medium dependent as well as strain dependent as bacteria have been shown to be diverse in their swarming behavior (Morris et al. 2011; Wang et al. 2004).

## Conclusion

While Pseudomonads act as strong competitors in homogeneous environments, in spatially structured environments they affect co-colonising *P. eucalypti* 299R positively. While this effect cannot be observed on a population scale during leaf colonization, we provide evidence that *P. eucalypti* 299R takes advantage of the presence of surfactant producing Pseudomonads during leaf colonization. This stresses the importance of surfactants produced by pseudomonads as public goods during leaf colonization and implies that cheating of *P. eucalypti* 299R and possibly other taxa may be the rule rather than the exception. Our study highlights another pivotal role of surfactants in the phyllosphere and the implications of surfactant production in this environment. Particularly in the context of preparing microbial inocula for the application on plants our findings provide additional traits, i.e. surfactant production and the ability for co-swarming, which should be considered during the formulation of products.

## Acknowledgements

The authors like to thank Sandra Hirsch for providing technical assistance.

## Contributions

MK, RS and MRE conceived the work. MK performed the experimental work. MK and MRE analyzed the data. RS and MRE supervised the work. LS provided material. MK and MRE wrote the initial draft of the manuscript with major input from all authors.

