## Supplemental data for "Hitching a ride in the phyllosphere: Surfactant production of *Pseudomonas* spp. causes co-swarming of *Pantoea eucalypti* 299R" (from another version of this preprint: version 1, 2023-11-08)

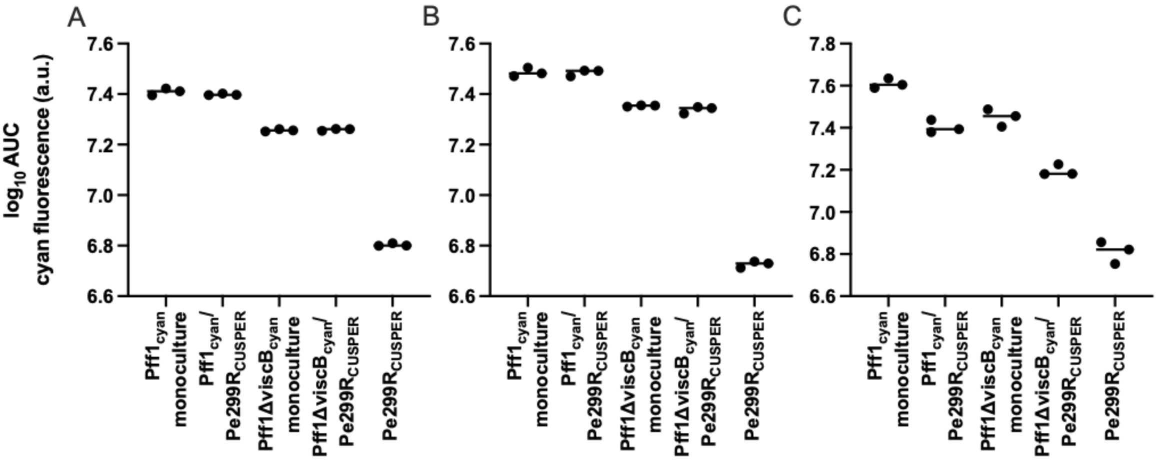


**Supplemental Figure 1** Impact of Pe299R$_{CUSPER}$ on growth of Pff1$_{cyan}$ and Pff1ΔviscB$_{cyan}$ in different media: A) M9 supplemented with glucose B) M9 supplemented with glucose, fructose and sorbitol and C) LB. Cyan fluorescence of Pe299R$_{CUSPER}$ Pff1$_{cyan}$ and Pff1ΔviscB$_{cyan}$ was tracked over time and the area under the curve was determined. Only the decrease of the cyan fluorescence signal during co-growth of Pe299R$_{CUSPER}$ and the Pseudomonads in rich medium is significantly different from the respective monocultures (unpaired t-test, $P < 0.001$).

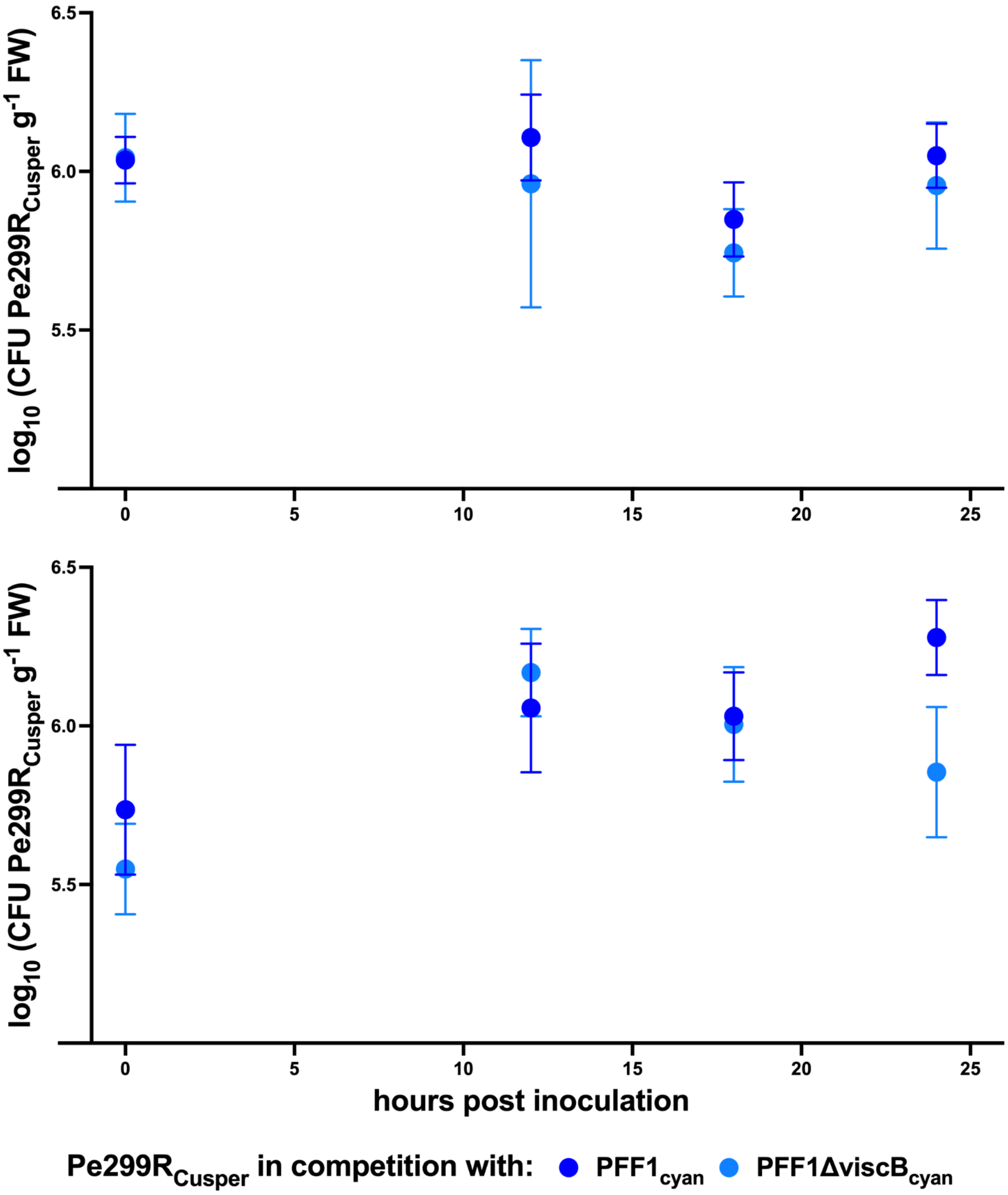


**Supplemental Figure 2** Growth of Pe299R$_{CUSPER}$ co-cultured with Pff1$_{cyan}$ and Pff1ΔviscB$_{cyan}$ *in planta*. No significant differences were detected on the population scale. For every treatment, four plants were harvested per time point, the experiment was performed three times independently. Two independent experiments are shown.

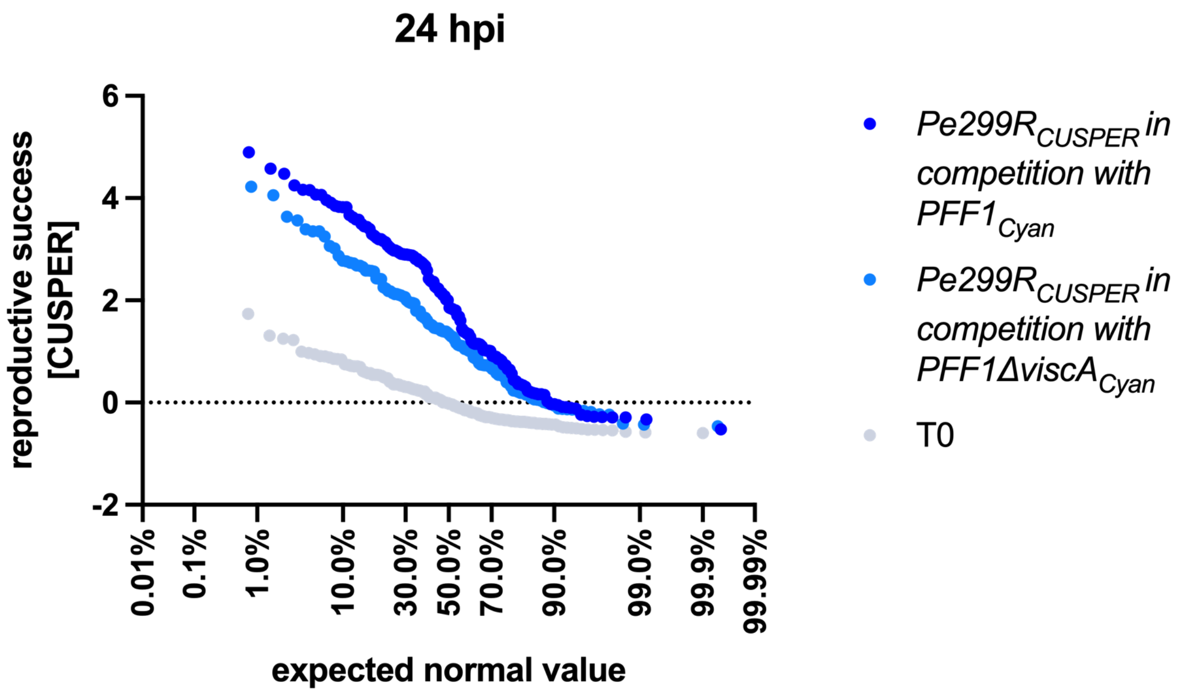


**Supplemental Figure 3** Reproductive success of individual Pe299R$_{CUSPER}$ cells during co-colonization of leaves with Pff1$_{cyan}$ or Pff1ΔviscB$_{cyan}$, third independent experiment. In gray, the respective T0 fluorescence intensity of the Pe299R$_{CUSPER}$ population is depicted. Every increase in reproductive success depicts a cell division relative to the T0 population. Every sample is pooled from the phyllosphere material of four individual plants..

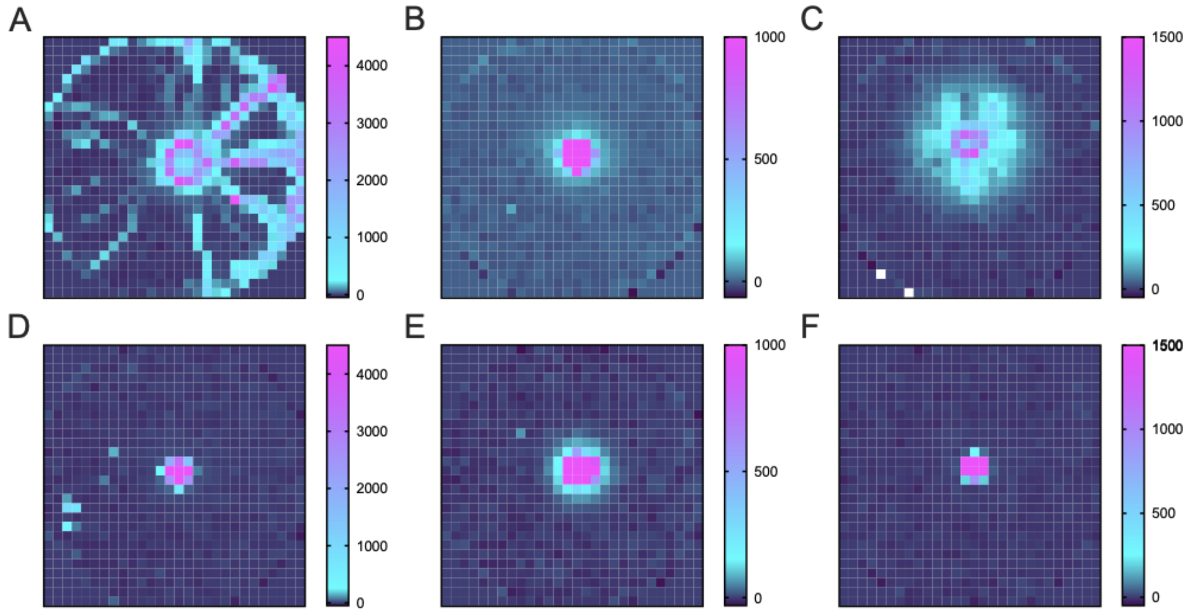


**Supplemental Figure 4** Pe299R$_{CUSPER}$ co-colonizing soft LB agar with Pff2, Pff3, Pff4 and their respective surfactant knockout mutants measured using a fluorescent plate reader (lower line). RFP emission depicts the distribution of Pe299R$_{CUSPER}$.

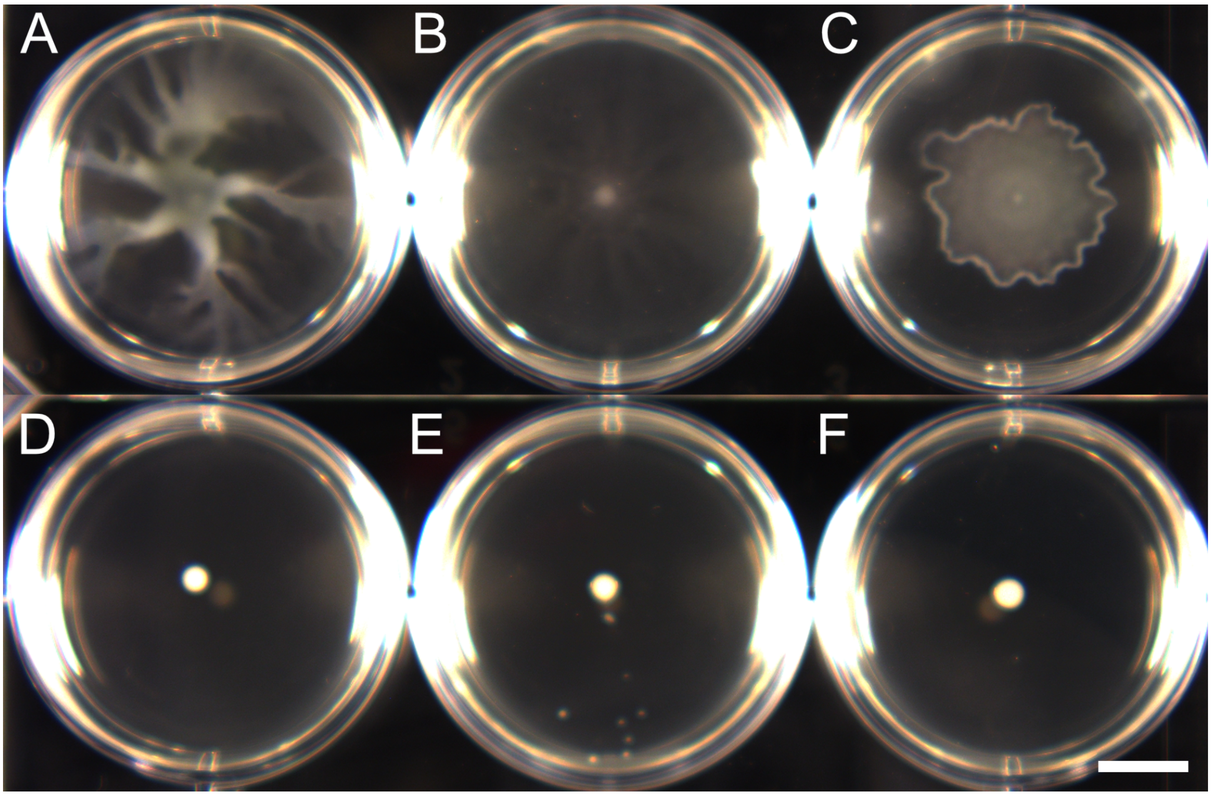


**Supplemental Figure 5** Macroscopical images of Pff2, Pff3, Pff4 and their knockout mutants on soft LB agar after 12 hours of growth. A-C) Wild type strains. D-F) Knockout mutants. Scale bar = 1 cm.
